# Translational evidence for RRM2 as a prognostic biomarker and therapeutic target in Ewing sarcoma

**DOI:** 10.1101/2021.03.04.433896

**Authors:** Shunya Ohmura, Aruna Marchetto, Martin F. Orth, Jing Li, Susanne Jabar, Andreas Ranft, Endrit Vinca, Katharina Ceranski, Martha J. Carreño-Gonzalez, Laura Romero-Pérez, Fabienne S. Wehweck, Julian Musa, Felix Bestvater, Maximilian M. L. Knott, Tilman L. B. Hölting, Wolfgang Hartmann, Uta Dirksen, Thomas Kirchner, Florencia Cidre-Aranaz, Thomas G. P. Grünewald

**Author notes:** correspondence Thomas G. P. Grünewald, MD, PhD Division Head, Division of Translational Pediatric Sarcoma Research (B410) German Cancer Research Center (DKFZ) & Hopp-Children’s Cancer Center (KiTZ) Im Neuenheimer Feld 280, 69210 Heidelberg, Germany.

## Abstract

**Purpose:** Ewing sarcoma (EwS) is a highly aggressive bone- or soft tissue-associated malignancy mostly affecting children, adolescents, and young adults. Although multimodal therapies have strongly improved patients’ overall survival over the past decades, the development of prognostic biomarkers for risk-based patient stratification and more effective therapies with less adverse effects is stagnating. Thus, new personalized medicine approaches are urgently required.

**Experimental design:** Gene expression data of EwS and normal tissues were crossed with survival data to identify highly overexpressed, prognostically relevant, and actionable potential targets. RNA-interference and dose-response assays as well as tissue-microarray analyses were carried out to explore the functional role and druggability of a prominent candidate gene *in vitro* and *in vivo*, and to validate its suitability as a prognostic biomarker.

**Results:** Employing a multilayered screening approach, we discover ribonucleotide reductase regulatory subunit M2 (RRM2) as a promising therapeutic target and prognostic biomarker in EwS. Through analysis of two independent EwS patient cohorts, we show that RRM2 mRNA and protein overexpression is associated with an aggressive clinical phenotype and poor patients’ overall survival. In agreement, *RRM2* silencing as well as pharmacological inhibition by the specific inhibitor triapine (3-AP) significantly reduces EwS growth *in vitro* and *in vivo*. Furthermore, we present evidence that pharmacological RRM2 inhibition by triapine can overcome chemoresistance against doxorubicin or gemcitabine, and synergize with cell cycle checkpoint inhibitors (CHEK1 or WEE1).

**Conclusions:** Based on the aggressive phenotype mediated by and the druggability of RRM2 our results provide a translational rationale for exploiting RRM2 as a novel therapeutic target in EwS and prompt further clinical investigations.

## INTRODUCTION

Ewing sarcoma (EwS) is an aggressive bone- or soft tissue-associated malignancy mostly affecting children, adolescents, and young adults (1). It is characterized by chromosomal translocations fusing members of the *FET* gene family (*EWSR1* or *FUS*) to variable *ETS* transcription factors, most commonly *FLI1* (85% of cases) (2, 3). The resulting fusion oncogene, *EWSR1-FLI1*, encodes an aberrant transcription factor that massively rewires the cellular transcriptome, which determines the malignant phenotype of EwS (1).

Clinically, EwS is a highly aggressive cancer with a high propensity for early hematological metastasis (4). The introduction of multiagent chemotherapy regimens including topoisomerase inhibitors (doxorubicin and etoposide), alkylating agents (cyclophosphamide and ifosfamide), and vinca alkaloids (vincristine) has remarkably improved patients’ overall survival since the 1970s (5). Despite this initial success, further therapeutic development for EwS has remained stagnant, especially for patients with metastatic or recurrent disease (6, 7). Besides, current multimodal therapies are frequently associated with early or late comorbidities in a substantial number of patients (8). Thus, more effective and in particular more specific treatment options are urgently required.

Although the pathognomonic fusion oncoprotein EWSR1-FLI1 would –in principle– constitute an ideal therapeutic target, its direct targeting by small molecule inhibitors or RNA interference remains challenging (9) due to the low immunogenicity of EWSR1-FLI1 encoded neoepitopes, the high and ubiquitous expression of its constituting genes, its intranuclear localization, and its lack of kinase or enzymatic activity (9).

To overcome these limitations, we have employed an alternative approach for investigating potential therapeutic targets by exploring putative downstream genes of EWSR1-FLI1 which are highly expressed in EwS compared to normal tissues and that promote the malignant phenotype of EwS. The former aspect may indicate a broad therapeutic window and less adverse effects, while the latter may minimize the risk for therapeutic escape. Prior scientificefforts have identified potential therapeutic targets for EwS treatment, which could offer avenues to personalized therapy (10–13). However, the discovery of directly clinically actionable downstream targets has remained in its infancy.

In the current study, we identified ribonucleotide reductase regulatory subunit M2 (RRM2) as a potential therapeutic target for a subset of EwS patients with high RRM2 expression. Our results demonstrate that high RRM2 expression confers an aggressive phenotype to EwS cells being associated with poor outcome, and that its genetic or pharmacologic inhibition reduces EwS growth in preclinical models, even in the setting of acquired chemoresistance. Finally, we show that inhibition of RRM2 by a specific inhibitor synergizes with cell cycle checkpoint inhibitors targeting CHEK1 or WEE1.

## RESULTS

### Ribonucleotide reductase regulatory subunit M2 (*RRM2*) is overexpressed in EwS, correlates with poor patient outcome, and constitutes a putative therapeutic target

To discover relatively specifically expressed, prognostically relevant, and druggable targets in EwS, we took advantage of publicly available ‘omics’ data and filtered them in a multi-step approach (**Fig. 1a**): First, we interrogated a curated gene expression dataset comprising 50 primary EwS and 929 samples from 71 normal tissue types to identify genes being overexpressed (min. log2 fold increase = 2) in EwS, which may offer a large therapeutic window. This analysis yielded 292 overexpressed genes (**Fig. 1b**, **Supplementary Table 1**). Second, we filtered for those genes whose overexpression was significantly negatively correlated with patients’ overall survival (*P*<0.05, Bonferroni-adjusted) in a dataset of matched gene expression and survival data of 166 EwS patients (11) that covered 280 of the 292 overexpressed genes (96%) (**Fig. 1c**). This filtering process identified 22 overexpressed genes with prognostic relevance (**Supplementary Table 1**). Third, among those 22 overexpressed and prognostically relevant genes, we focused on druggable targets possessing kinase or other enzymatic functions for which specific inhibitors and their pharmacokinetic data were already available, but were still not (pre)clinically tested in EwS. This survey identified ribonucleotide reductase regulatory subunit M2 (*RRM2*) as the single putative target that fulfilled our above-mentioned selection criteria, and that exhibited a prominently negative association with patients’ overall survival (**Fig. 1d**). RRM2 is the regulatory subunit of the ribonucleotide reductase (RNR), which is a heterotetrameric holoenzyme catalyzing the rate-limiting *de novo* conversion of ribonucleoside diphosphates to deoxyribonucleoside diphosphates (14). RNR is composed of two subunits, ribonucleotide reductase catalytic subunit M1 (RRM1) and either RRM2 or ribonucleotide reductase regulatory TP53 inducible subunit M2 (RRM2B) (14). While RRM1:RRM2 mainly contributes to DNA synthesis and DNA repair during S-phase, RRM1:RRM2B is responsible for DNA repair in quiescent cells and for mitochondrial DNA synthesis or repair (14). Notably, *RRM2B* is neither overexpressed in EwS nor negatively correlates with patient outcome (**Supplementary Figs. 1a,b**), and *RRM1* is far less overexpressed in EwS compared to *RRM2* (**Supplementary Figs. 1a,b**). Thus, we henceforth focused on *RRM2* in subsequent functional experiments.

**Fig. 1.**
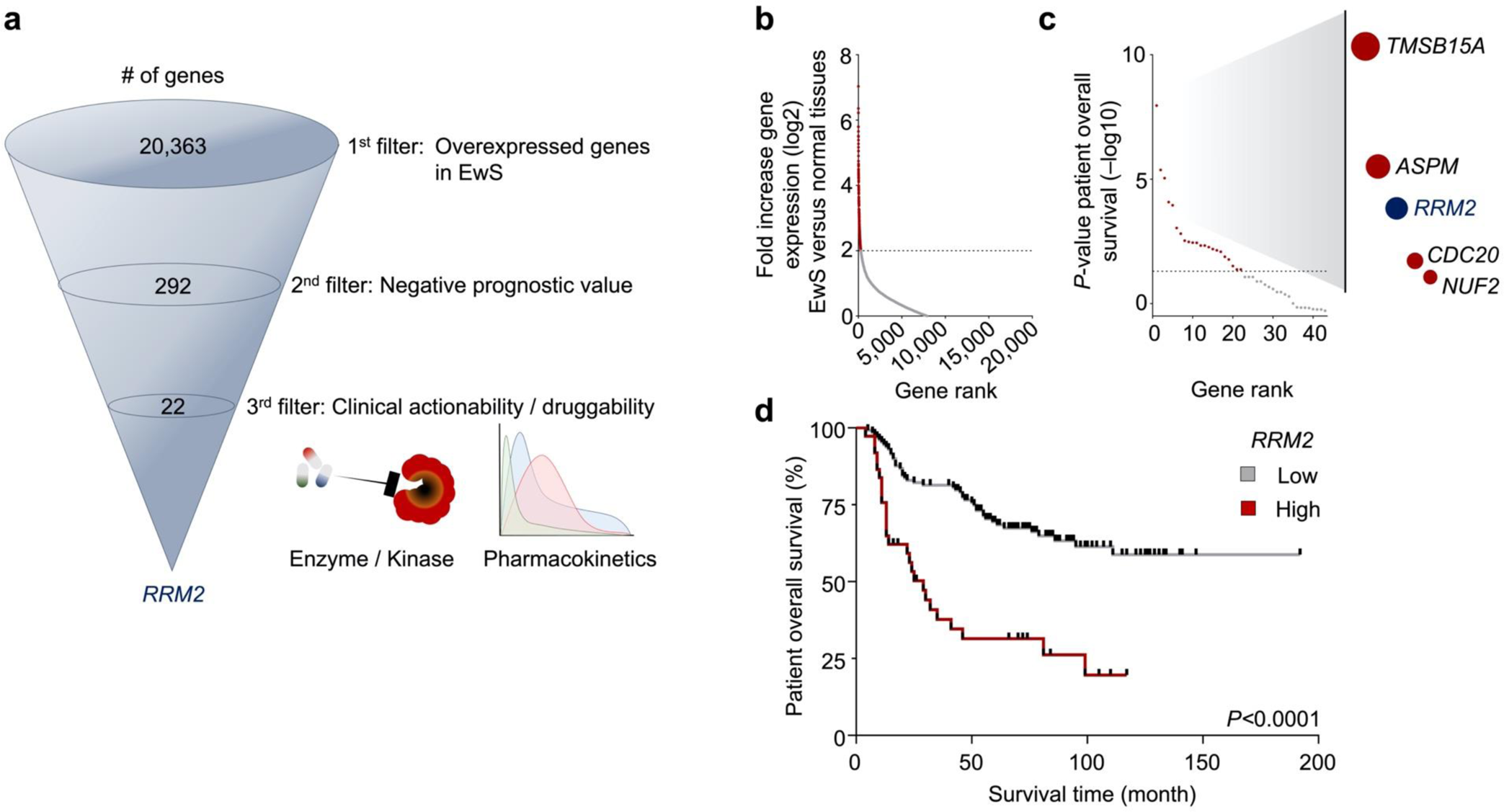
*RRM2* is highly overexpressed in EwS, correlates with poor patient outcome, and constitutes a putative therapeutic target. **a)** Schematic description of the filtering process for identification of therapeutically relevant target candidates. **b)** Analysis of *RRM2* mRNA expression levels in 50 EwS primary tumors compared to 929 normal tissues samples from 71 tissue types. Data are shown as log2 fold increase normalized to expression values of normal tissues. The dotted line indicates the cut-off value of 2 for candidate selection. **c)** Analysis of overall survival time of 166 EwS patients stratified for candidate gene expression. *P*-values (–log10) were determined in Kaplan-Meier analyses using a Mantel-Haenszel test (Bonferroni-adjusted for multiple testing). The dotted line indicates a significance value of 1.3. **d)** Kaplan-Meier survival analysis of 166 EwS patients stratified by the 78th percentile *RRM2* expression. *P*-value determined by log-rank test.

### RRM2 overexpression is correlated with an aggressive tumor phenotype in EwS

Prior reports focusing mainly on three EwS cell lines suggested that RRM2 may contribute to the proliferative phenotype of EwS (15–19). However, its role in primary EwS tumors remained unclear. To gain first insights into the biological function of *RRM2* in EwS tumors, we carried out gene ontology (GO) enrichment analysis of *RRM2* co-expressed genes in 166 EwS tumors, which revealed that high *RRM2* expression is closely correlated with cell cycle- (or cell division-) associated gene expression signatures (**Fig. 2a**), suggesting that high *RRM2* expression may contribute to an aggressive clinical course by promoting tumor growth. To test this hypothesis, and to validate the prognostic role of RRM2 in a second cohort and on the protein level, we analyzed the potential association between RRM2 protein expression levels, known clinicopathological prognostic markers, and clinical outcomes in tissue microarrays (TMA) from EwS tumors of 122 patients (**Table 1**, **Fig. 2b**). In agreement with our findings at the mRNA level (**Fig. 1d**), high RRM2 protein expression as detected by immunohistochemistry (IHC) was significantly (*P*=0.0095) associated with poor overall survival (**Fig. 2c**). In addition, we performed correspondence analysis of each cohort individually as well as a joint analysis compiling both cohorts (after exclusion of 6 samples (3.6 %) from the mRNA-cohort that were in overlap with the TMA cohort), which revealed that high RRM2 expression was significantly associated with metastatic disease at diagnosis (*P*=0.0004) and occurrence of metastatic and/or local relapse (*P*=0.0095; information only available for the TMA cohort) (**Table 2**), supporting that high RRM2 expression promotes an aggressive phenotype.

**Fig. 2.**
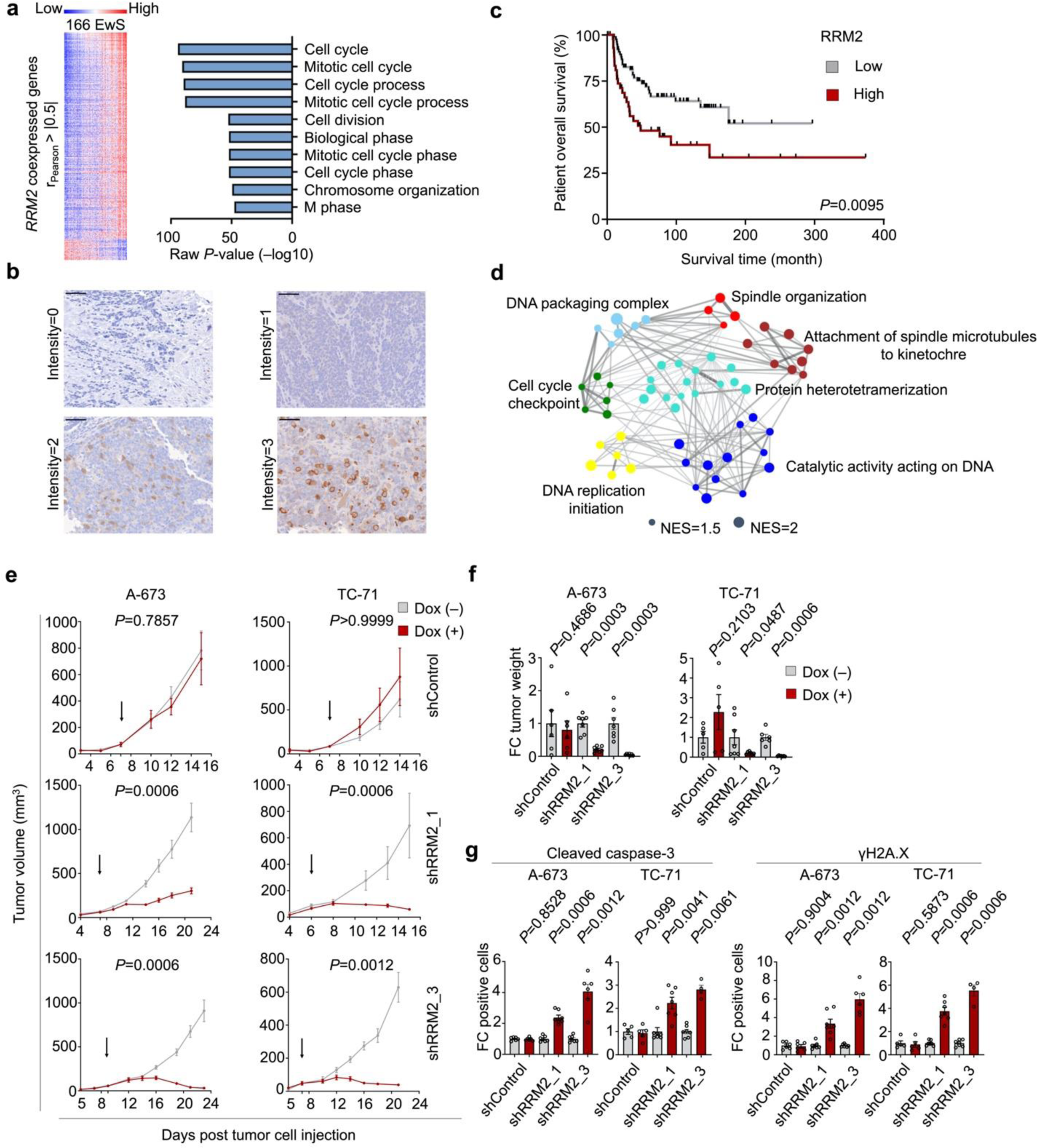
RRM2 overexpression is correlated with an aggressive tumor phenotype in EwS. **a)** Left: Heat map for gene expression which is positively or negatively correlated with *RRM2* expression in 166 EwS. Right: Gene ontology (GO) enrichment analysis of *RRM2* and its co-expressed genes derived from gene expression data sets of 166 EwS tumors. Pearson correlation coefficients between *RRM2* and other genes were determined, of which those with |r_Pearson_| > 0.5 were further analyzed by GO enrichment analysis. **b)** Representative images of immunohistochemical RRM2 staining. Scale bar = 50 µm. **c)** Kaplan-Meier survival analysis of 122 EwS patients stratified by RRM2 protein expression (low IRS≤2, high IRS >2). *P*-values were determined by log-rank test. **d)** WGCNA of downregulated genes upon *RRM2* silencing in A-673 and ES-7 cells harboring Dox-inducible shRRM2 constructs. NES, normalized enrichment score. **e)** Analysis of tumor growth of EwS cell lines A-673 and TC-71 harboring Dox-inducible shRRM2 constructs or non-targeting shRNA (shControl) xenografted in NSG mice. Once tumors were palpable, animals were randomized in Dox (+) or Dox (–) group. Tumor growth on time course and **f)** Tumor weight at the experimental endpoint. Arrows indicate treatment start. Values are normalized to shControl. Horizontal bars represent means and whiskers SEM. FC, fold change. *P*-values were calculated at the experimental endpoint with two-sided (tumor growth) or one-sided (tumor weight) Mann-Whitney test. **g)** Representative micrographs of xenografts immunohistochemically stained for RRM2, cleaved caspase-3 (CC3) or γH2A.X (scale bar=250 µm, 50 µm, 250 µm, respectively). **h)** quantification of positive cells for cleaved caspase-3 (CC3) (left) and γH2A.X (right). Values were normalized to shControl. Horizontal bars represent means and whiskers SEM. FC, fold change. *P*-values were calculated at the experimental endpoint using a two-sided Mann-Whitney test.

**Table 1.** Clinicopathological characteristics of EwS patients for the mRNA (discovery) and TMA-cohort (validation).

|  | n=166 | n=122 |
| --- | --- | --- |
| Variable | Discovery-cohort (mRNA) n (%) | Validation-cohort (TMA) n (%) |
| Sex |  |  |
| Male | 94 (56.6) | 75 (64.1) |
| Female | 72 (43.4) | 42 (35.9) |
| Missing |  | 5 |
| Age at diagnosis |  |  |
| <15 years | 83 (50) | 37 (31.6) |
| ≥15 years | 83 (50) | 80 (68.4) |
| Missing |  | 5 |
| Metastasis at diagnosis |  |  |
| Absent | 107 (86.6) | 84 (71.8) |
| Present | 18 (14.4) | 33 (28.2) |
| Missing | 41 | 5 |
| Site |  |  |
| Axial | 31 (56.4) | 58 (50.9) |
| Non-axial | 24 (43.6) | 56 (49.1) |
| Missing | 111 | 8 |
| Tumour volume |  |  |
| <200 ml | NA | 47 (56.6) |
| ≥200 ml | NA | 36 (43.4) |
| Missing |  | 39 |
| Histological response |  |  |
| Good | NA | 46 (79.3) |
| Poor | NA | 12 (20.7) |
| Missing |  | 64 |
| Status |  |  |
| Alive | 95 (57.2) | 72 (59.0) |
| Dead | 71 (42.8) | 50 (41.0) |
| Missing |  |  |
| RRM2 expression |  |  |
| High | 37 (22.3) | 43 (35.2) |
| Low | 129 (77.7) | 79 (64.8) |
| NA = not available. |  |  |

**Table 2.**
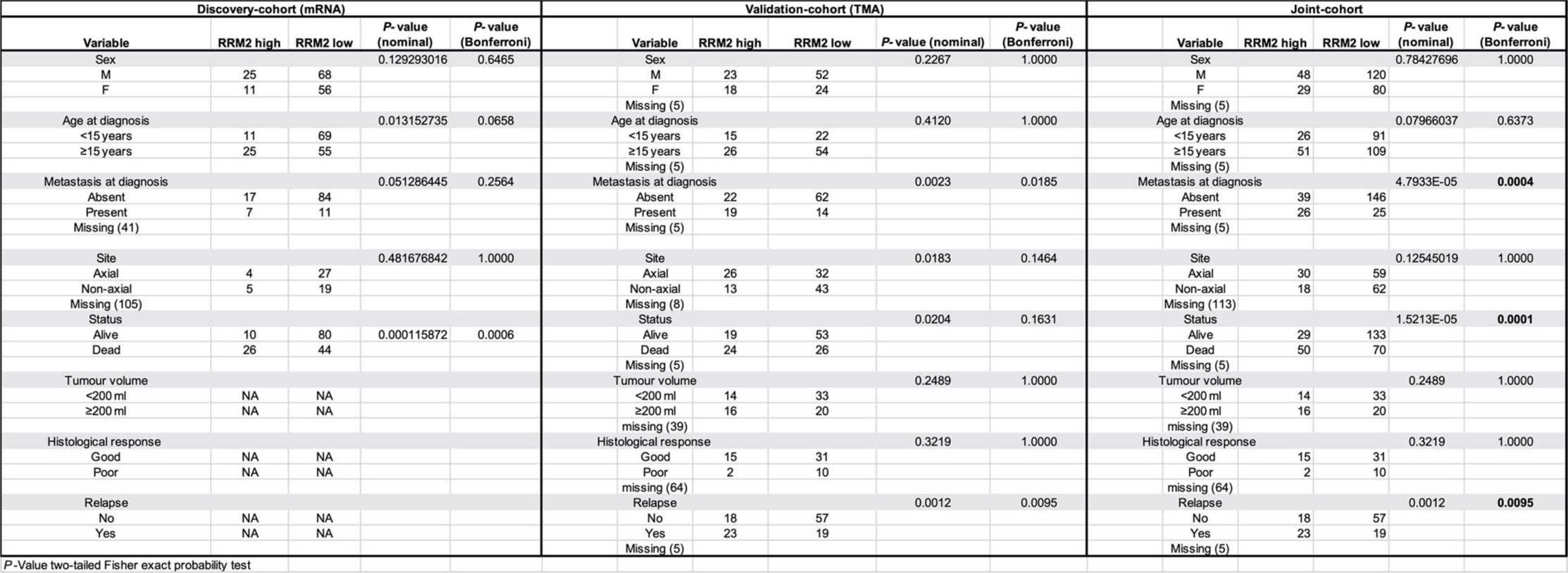
Multivariate analysis for RRM2 expression and clinical parameters in the mRNA (discovery), TMA-cohort (validation) and joint-cohort.

In agreement with these clinical associations and previous data (15–19), doxycycline (Dox)-inducible RNA interference (RNAi)-mediated *RRM2* silencing inhibited proliferation and clonogenic growth of A-673, ES7 and TC-71 EwS cell lines, and induced cell death *in vitro* (**Supplementary Figs. 2a,b**). In line with these functional experiments, gene expression profiling of *RRM2*-silenced EwS cell lines (A-673, ES7) by Affymetrix Clariom D microarrays followed by gene set enrichment analysis (GSEA) and weighted gene co-expression network analysis (WGCNA) demonstrated a strong downregulation of cell cycle and proliferation-associated gene signatures (**Fig. 2d**). Similarly, conditional knockdown of RRM2 significantly reduced tumor growth of two xenografted EwS cell lines (A-673, TC-71) *in vivo*, which was not observed for xenografts harboring an inducible negative control shRNA (**Figs. 2e,f**). This antineoplastic effect was accompanied by increased apoptotic rates and DNA damage, as assessed by IHC for cleaved caspase 3 (CC3) and γH2A.X, respectively (**Fig. 2g**, **Supplementary Fig. 2c**). Collectively, these results provided evidence that RRM2 promotes tumor growth and aggressive behavior of EwS, and further indicated that it may constitute a promising therapeutic target.

### Triapine inhibits EwS growth *in vitro* and *in vivo*

Since RRM2 appeared as a therapeutic target in EwS, we aimed at pharmacological blocking of its function. Generally, the activity of RNR can be blocked by inhibiting RRM1 using a RRM1-specific inhibitor such as gemcitabine, or by RRM2-specific inhibitors such as hydroxyurea or a more potent drug triapine (alias 3-AP) (14, 20). Gemcitabine is used in conjunction with other antitumor agents such as taxanes for palliative treatments of EwS patients (21), and earlier reports have shown antineoplastic effects of gemcitabine in combination with CHEK1-inhibitors in short-term preclinical models (19). Yet, many EwS patients seem to rapidly develop a relative resistance toward gemcitabine (21). In support of this observation, we found that long-term treatment of the EwS cell lines with ascending doses of either doxorubicin (A-673, ES7, EW-7, TC-71), gemcitabine (A-673, ES7, TC-71) or triapine (A-673) led to acquisition of relative resistance phenotypes *in vitro* (**Supplementary Fig. 3a**). Through analysis on development rates of drug resistance in A-673 we noted a relatively fast and strong increase of the relative resistance toward gemcitabine (>2,000-fold increase in IC_50_ within ∼6 weeks), the same cells exhibited a much lower increase in IC50 after a substantially longer period of treatment time for doxorubicin (∼4-fold increase in ∼28 weeks) and triapine (∼7-fold increase in ∼20 weeks) (**Supplementary Fig. 3b**), further suggesting that gemcitabine has limited potential for clinical treatment with curative intent.

Thus, we tested the antineoplastic effects of triapine *in vitro* and *in vivo*. Dose-response assays revealed that EwS cell lines were very sensitive toward pharmacological RRM2 inhibition by triapine compared to osteosarcoma cell lines and non-transformed EwS patient-derived mesenchymal stem cells (mean IC50 values 0.35, 1.63, 101.63 µM, respectively) (**Fig. 3a**). Likewise, triapine treatment significantly reduced clonogenic growth of EwS cell lines at clinically relevant doses (22, 23) (**Fig. 3b**). Interestingly, doxorubicin or gemcitabine resistant EwS cells (designated EwS/DR or EwS/GR, respectively) still retained their triapine sensitivity (**Fig. 3c**), suggesting a therapeutic potential of triapine for EwS refractory toward conventional chemotherapy.

**Fig. 3.**
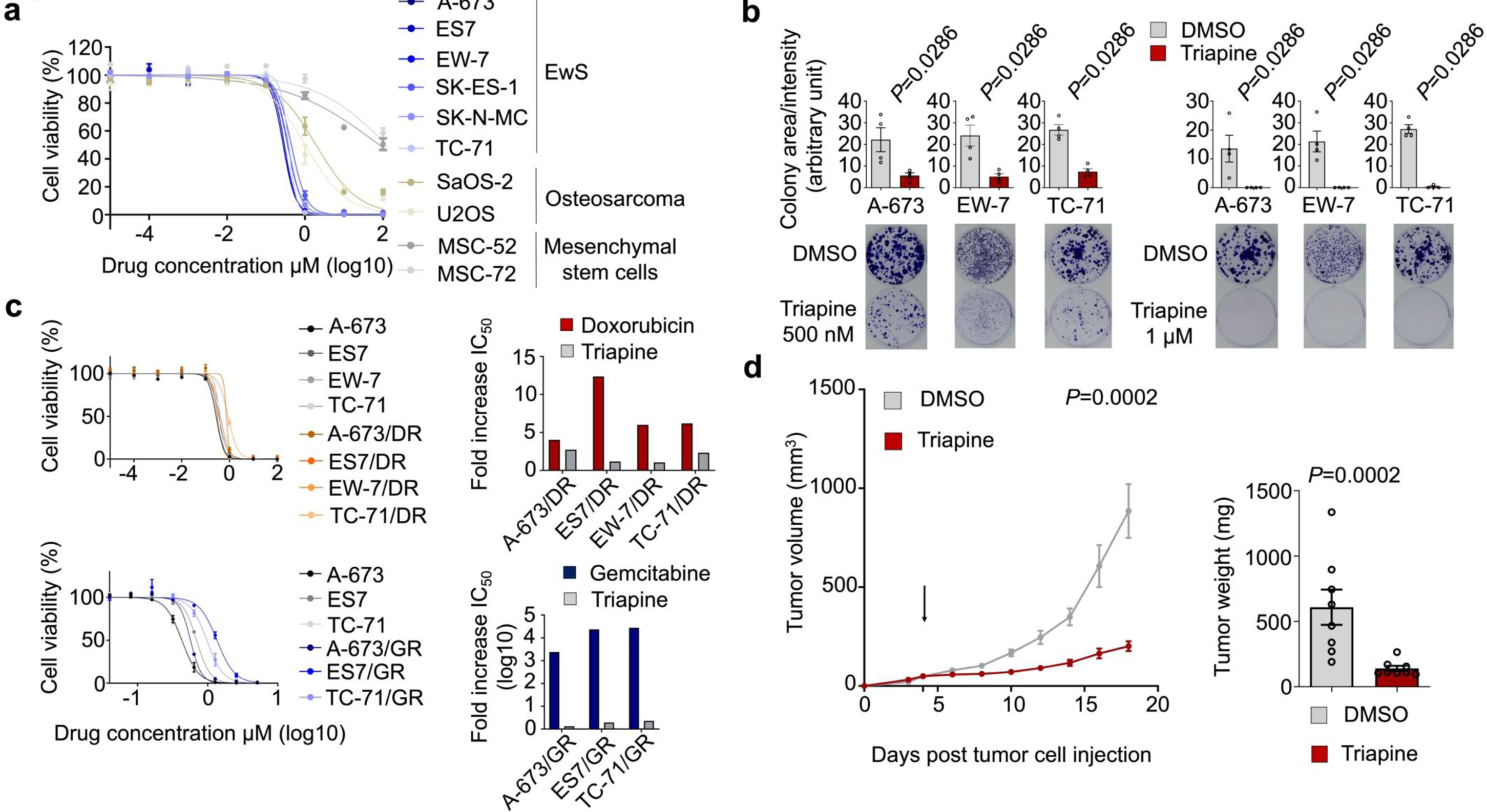
Triapine inhibits EwS growth *in vitro* and *in vivo*. **a)** Dose response analysis of triapine in EwS, osteosarcoma and mesenchymal stem cells. **b)** Analysis of clonogenic growth of A-673, TC-71, and EW-7 EwS cells upon triapine treatment. **c)** Left: Dose-response analysis of triapine in chemoresistant EwS cells (A-673/DR or A-673/GR). Right: magnitudes of doxorubicin or gemcitabine resistance shown by fold increase in IC50 compared to those of parental cells. **d)** Analysis of tumor growth upon triapine treatment in A-673 cell line *in vivo*. Tumor growth (left) and tumor weight at the experimental endpoint (right) of NSG mice xenografted with A-673 EwS cells upon treatment with triapine. Once tumors reached 5 mm in average diameter, animals were randomized in treatment group (30 mg/kg i.p.) or control group (DMSO) (n=8 animals per group). The arrow indicates treatment start. Horizontal bars represent means and whiskers SEM. *P*-values were calculated at the experimental endpoint with two-sided (tumor growth) or one-sided (tumor weight) Mann-Whitney test.

Next, we assessed the antineoplastic effects of triapine and its possible adverse effects *in vivo*. To this end, we xenografted A-673 cells subcutaneously in immunocompromised NOD/SCID/gamma (NSG) mice and monitored tumor growth over time during triapine treatment (30 mg/kg triapine i.p. every second day once tumors reached ∼5 mm in average diameter). Strikingly, we observed a significant reduction of tumor growth compared to controls (DMSO) (**Figs. 3d**). Although triapine treatment was accompanied by a statistically significant weight loss of ∼10% at the experimental endpoint (**Supplementary Fig. 3c**), which is in line with the known adverse effect profile of triapine shown by several clinical studies (24, 25), neither other adverse effects nor morphological changes of inner organs were observed including the gastrointestinal tract as assessed by histological analysis (**Supplementary Fig. 3d**). Together these preclinical data indicate that triapine could constitute a novel effective agent not only for chemotherapy-naive but also for chemotherapy-refractory EwS. To mitigate its adverse effects, however, a combinatorial approach would be desirable.

### Triapine synergizes with cell cycle checkpoint inhibitors in EwS *in vitro*

Since in the clinical setting most cancers rapidly acquire resistance phenotypes against monotherapies with chemotherapeutics or small molecule inhibitors (26, 27), and considering the adverse effects, we next explored effective drug combinations with triapine. Based on known functions of RRM2 in DNA synthesis and DNA repair (14) we hypothesized that pharmacological RRM2 inhibition with triapine could synergize with the antineoplastic effects of chemotherapeutics, such as doxorubicin, etoposide, and vincristine, which are routinely employed in EwS treatment protocols (4), or inhibition of DNA repair-associated targets, such as poly ADP-ribose polymerase (PARP) (28, 29). Surprisingly, we did not observe any marked synergistic effects but rather antagonistic effects (Supp. **Fig. 4a**). Next, to systematically identify further rational combinatorial interventions, we analyzed gene expression profiles of EwS cells upon *RRM2* silencing and pharmacological inhibition by triapine. By integrating the expression profiles of two EwS cell lines A-673 and ES7 upon *RRM2* silencing and triapine treatment we found 263 commonly up- and downregulated genes (**Supplementary Table 2**). Gene ontology (GO) enrichment analysis demonstrated significant enrichment for cell cycle-associated processes, especially regulation of mitotic cell cycle-associated genes (**Fig. 4a**), which is in line with the observation that RRM2 inhibition caused G1/S-phase cell cycle arrest (30, 31). Thus, we reasoned RRM2 may synergize with checkpoint inhibitors targeting CHEK1 (checkpoint kinase 1) or WEE1 (WEE1 G2 checkpoint kinase) that were highly significantly (*P*<0.0001) co-expressed with *RRM2* in the microarray gene expression data set of 166 EwS tumors (**Fig. 4b**). In support of this hypothesis, other RNR inhibitors, such as gemcitabine or hydroxyurea, were reported to synergize with CHEK1 or WEE1 cell cycle checkpoint inhibitors (32, 33). In drug combination assays, in the four EwS cell lines tested, we observed a strong synergism between triapine and the CHEK1 inhibitor (CCT245737) or the WEE1 inhibitor (MK-1775), both of which are currently under clinical investigation (ClinicalTrials.gov Identifier: NCT02797964, NCT03668340) (Figs. 4c, d, **supplementary Fig. 4b**).

**Fig. 4.**
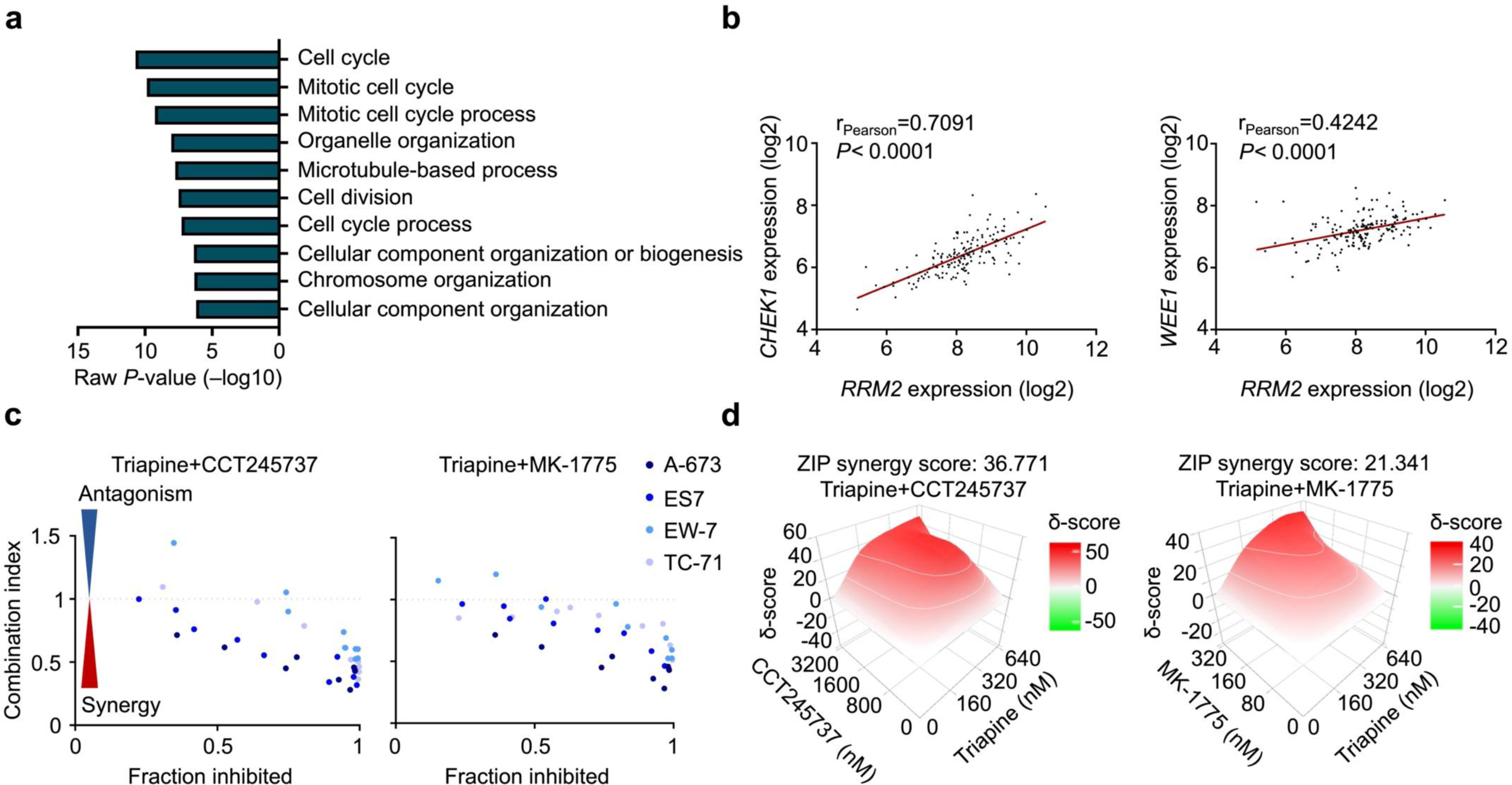
Triapine synergizes with cell cycle checkpoint inhibitors in EwS *in vitro*. **a)** Integrative Gene Ontology (GO) enrichment analysis of gene expression microarray data generated in A-673 and ES7 cells after *RRM2* silencing or pharmacological RRM2 inhibition by triapine (corresponding IC50 of 0.44 µM or 0.65 µM, respectively). **b)** Correlation of gene expression between *RRM2* and *CHEK1* or *WEE1* in 166 EwS. Each dot represents an individual expression value. Solid red lines indicate a trend line created by a simple linear regression. *P*-values were calculated by a two-tailed t-test. **c)** Drug interaction and combination efficiency analysis between triapine and CHEK1 inhibitor (CCT245737) or WEE1 inhibitor (MK-1775) in 4 EwS cell lines (A-673, ES7, EW-7, TC-71) assessed by combination index. CI value < 1 indicative of synergistic, CI = 1 additive, and CI > 1 antagonistic **d)** Drug interaction and combination efficiency estimation between triapine and CHEK1 inhibitor (CCT245737) or WEE1 inhibitor (MK-1775) in A-673 EwS cell line assessed by SynergyFinder 2.0. ZIP synergy score > 10, likely to be synergistic; between -10 and 10, likely to be additive; < –10, likely to be antagonistic.

Overall, these results provide a rationale for therapeutic combination of triapine with cell cycle checkpoints inhibitors for EwS treatment.

## DISCUSSION

Through our multilayered screening approach, we demonstrated that 1) *RRM2* is highly expressed in EwS compared to normal tissues, 2) RRM2 mRNA and protein overexpression is correlated with clinically aggressive phenotype, 3) RRM2 inhibition by gene silencing with RNA interference or pharmacological inhibition by triapine reduces EwS growth *in vitro* and *in vivo*, 4) pharmacological RRM2 inhibition by triapine is effective even in doxorubicin or gemcitabine resistant EwS cell lines, and 5) triapine synergizes with CHEK1 or WEE1 inhibitors. Our results provide a rationale for therapeutically targeting RRM2 by triapine and its combination with CHEK1 or WEE1 inhibitors not only for chemotherapy-naive but also for chemotherapy-refractory EwS.

Earlier studies have shown that RNR could constitute a potential therapeutic target in EwS (17), and demonstrated that gemcitabine can synergize with cell cycle checkpoint inhibitors, such as CHEK1 in preclinical models (19). In our study, we extended this research and also pointed out issues of chemoresistance. Gemcitabine has been clinically used mainly for treatment of otherwise chemotherapy-refractory EwS (21). Unfortunately, gemcitabine is only effective in a subset of EwS patients and in most patients only for a short period of time (21, 34). These clinical observations align with our findings of a rapidly acquired resistance toward gemcitabine in the A-673 EwS cell line. Generally, chemoresistance against current standard chemotherapeutics represents one of the major clinical obstacles (26, 27). The results of our preclinical models suggest that triapine could be used as a novel candidate drug to overcome not only doxorubicin-, but also gemcitabine-resistance in EwS, which may warrant a validation in future clinical trials.

Since single-agent-therapy is relatively rapidly bypassed by cancer cells, which we could also observe for triapine in our *in vitro* models, we explored an effective drug combination to mitigate triapine-resistance and potential adverse effects. Although earlier studies demonstrated a synergistic effect of triapine with chemotherapeutics or PARP inhibitors in other cancer types (35, 36), we did not observe marked synergy with these drug combinations in EwS cells, which may indicate tumor-specific effects with combinatorial drug applications. Importantly, similar to a very recent report in neuroblastoma (37), we noted a strong synergistic effect of triapine with CHEK1 or WEE1 inhibitors in EwS cells.

The major issue on combining RNR inhibitors with cell cycle checkpoint inhibitors is adverse effects (38), which could be caused by extended DNA damage. For example, gemcitabine combined with CHEK1 inhibitor (GDC-0575) appears to cause strong dose-dependent myelosuppression (32, 38). Gemcitabine binds covalently to RRM1 after its intracellular phosphorylation by deoxycytidine kinase converting it to difluorodeoxycytidine diphosphate, which results in irreversible inhibition of RNR. In contrast, triapine seems to inhibit RNR activity only transiently by inhibiting the generation of tyrosyl radicals at RRM2 subunits (14). Given the much larger fold increase of *RRM2* expression in EwS compared to normal tissues and its transient inhibitory nature, it is tempting to speculate that triapine combined with cell cycle checkpoint inhibitors may offer a more specific and controllable treatment option than gemcitabine as recently demonstrated by a drug combination of hydroxyurea and CHEK1 inhibitor GDC-0575 (32) – an aspect being subject to ongoing research. Yet, it should be noted that triapine treatment can also cause myelosuppression and a unique adverse effect i.e. transient meth-hemoglobinemia by short-term triapine infusion (∼30% of patients) (39, 40), which can be explained by its iron-chelating function (41). Interestingly, prolonged infusion of triapine can circumvent this unique toxicity (23). Alternatively, an oral formulation or combination therapies could potentially mitigate this toxicity (22).

The regulatory mechanism(s) controlling RRM2 expression in EwS remain largely uncharacterized. A very recent study proposed that RRM2 may be translationally regulated by eukaryotic translation initiation factor 4E binding protein 1 (EIF4EBP1, also known as 4E-BP1) in *in vitro* models (16). While we noted that RRM2 protein is heterogeneously overexpressed across EwS tumors, we also observed its heterogeneous transcription levels in primary EwS (**Supplementary Fig. 5a**), which suggests a more complex and multi-layered regulatory mechanism of *RRM2* expression in EwS. In this context, we did neither observe a significant correlation of *RRM2* expression with copy number alterations at the *RRM2* locus nor with methylation status of the promoter regions in primary EwS tumors (**Supplementary Figs. 5b,c**). Yet, since *RRM2* has been shown to be transcriptionally upregulated by *E2F1* (42), which can be in turn upregulated by EWSR1-FLI1 (43), we assume an indirect regulatory mechanism of RRM2 overexpression by EWSR1-FLI1 in EwS, which should be further investigated in future studies.

Clinical associations of *RRM2* overexpression with inferior chemotherapy response has been shown in several tumor types including EwS (14, 44). In line with our observation in TMA analysis, Schaefer *et al.* showed by comparative transcriptome analysis of primary EwS tumors in chemotherapy responders/non-responders that high *RRM2* expression is associated with a refractory phenotype to neoadjuvant standard chemotherapy for EwS (44). Together these findings further support a role of RRM2 as a potential biomarker to identify a subgroup of EwS patients with higher risk for a clinically aggressive phenotype, which might in turn benefit in particular from co-treatment with the RRM2-specific inhibitor triapine. In this regard, it is interesting to note that triapine was still effective in our otherwise chemotherapy-resistant EwS models (**Fig. 3c**).

In summary, we demonstrated that RRM2 can be a promising therapeutic target not only for chemotherapy-naive but for chemotherapy-refractory EwS. Combination of triapine with cell cycle checkpoint inhibitors such as for CHEK1 or WEE1 could offer more effective therapy with less adverse effects, which we are addressing in our ongoing research.

## ACKNOWLEDGEMENTS

We are grateful to M. Melz for constructing TMAs, to S. Stein and F. Zahnow for experimental assistance, and to A. Heier and A. Sendelhofert for technical assistance in establishing immunohistochemical staining procedures. This work was mainly supported by a grant from the Deutsche Forschungsgemeinschaft (DFG-391665916 to S.O.). In addition, the laboratory of T.G.P.G. is supported the Matthias-Lackas Foundation, the Dr. Leopold and Carmen Ellinger Foundation, the Dr. Rolf M. Schwiete Foundation, the German Cancer Aid (DKH-70112257; DKH-70114111), the Gert and Susanna Mayer Foundation, the SMARCB1-association, and the Barbara and Wilfried Mohr foundation. The working group of U.D. (S.J. and A.R) is supported by the German Cancer Aid (DKH-70113419, DKH-70112418), the Gert and Susanna Mayer Foundation, and the Barbara and Hubertus Trettner Foundation. T.L.B.H. was supported by a Mildred-Scheel scholarship from the German Cancer Aid.

## AUTHOR CONTRIBUTIONS

S.O. and T.G.P.G. conceived the study, wrote the manuscript and designed the figures and tables. S.O., A.M., and E.V. performed *in vitro* experiments. S.O. performed *in vivo* experiments, and F.C.A. helped in coordination of *in vivo* experiments. S.O., M.F.O., K.C., M.J.C.G., F.C.A., J.L., and T.G.P.G. performed bioinformatic and statistical analyses. S.O. and F.W. scored tissue-microarrays. S.J., A.R., U.D., and W.H. provided gene expression data, tissue-microarrays and helped in statistical analysis of clinical data. M.F.O., J.L., F.C.A., L.R.P., M.M.L.K., T.L.B.H., J.M. contributed to experimental procedures. F.B. and T.K. provided laboratory infra-structure and histological guidance. T.G.P.G. supervised the study and data analysis. All authors read and approved the final manuscript.

## MATERIAL AND METHODS

### Cell lines and cell culture conditions

Human HEK293T cells and the human Ewing sarcoma (EwS) cell line A-673 were purchased from the American Type Culture Collection (ATCC). Human EwS cell lines SK-ES-1, and SK-N-MC, as well as the human osteosarcoma cell lines SaOS-2 and U2OS were provided by the German Collection of Microorganisms and Cell Cultures (DSMZ). Human EwS cell line TC-71 was kindly gifted by the Children’s Oncology Group (COG). Human EwS cell lines ES7 and EW-7 were provided by O. Delattre (Institute Curie, Paris). The human mesenchymal stem cell lines MSC-52 and MSC-72 originated from tumor-free bone marrow of EwS patients were provided by U. Dirksen (Essen, Germany). All cell lines except for MSC-52 and MSC-72 were maintained in RPMI-1640 media with stable glutamine (Biochrom, Germany) supplemented with 10% tetracycline-free fetal bovine serum (Sigma-Aldrich, Germany), 100 U/ml penicillin and 100 µg/ml streptomycin (Merck, Germany) in a humidified incubator with 5% CO_2_ at 37°C. MSC-52 and MSC-72 were maintained in Alpha MEM (Biochrom, Germany) supplemented with 10% tetracycline-free fetal bovine serum, 100 U/ml penicillin and 100 µg/ml streptomycin, 1% L-glutamine (Thermo Fisher Scientific) and 2 ng/ml recombinant human FGF-Basic (Thermo Fisher Scientific) in a humidified incubator with 5% CO_2_ at 37°C. All cell lines were routinely tested for the absent of mycoplasma contamination by nested-PCR. STR-profiling was regularly performed to assess cellular identity.

### Chemical compounds

CCT245737, doxorubicin, etoposide, gemcitabine, MK-1775, niraparib, olaparib, triapine (3-AP), and vincristine were purchased from Selleckchem, resolved in DMSO and stored at –80°C.

### Assessment of mRNA expression with quantitative real-time polymerase chain reaction (qRT-PCR)

Total RNA was isolated using NucleoSpin RNA kit according to the manufacturer’s instruction (Macherey-Nagel, Germany). One µg of total RNA was reverse-transcribed with High-Capacity cDNA Reverse Transcription Kit (Applied Biosystems, USA). cDNA was diluted at 1:10 in ddH_2_O and stored at -20°C. For qRT-PCR 6.75 µl cDNA was mixed with 7.5 µl SYBR green Master Mix (Applied Biosystems), 0.75 µl forward and reverse primer (10 µM) in a total volume of 15 µl. qRT-PCR reactions were performed with a BioRad CFX Connect instrument and the following thermal cycles; heat activation at 95°C for 2 min, denaturation at 95°C for 10 sec, annealing and elongation at 60°C for 20 sec (50 cycles), final denaturation at 95°C for 30 sec. Data was analyzed by BioRad CFX Manager 3.1 software. For assessment of mRNA expression levels, the 2−ΔΔCt method (45) was employed normalized to the housekeeping gene *RPLP0*. Oligonucleotides were purchased from MWG Eurofins Genomics (Ebersberg, Germany). Sequences are listed in **Supplementary Table 3**.

### Doxycycline (Dox)-inducible target knockdown by RNA interference with short hairpin RNA (shRNA)

The Tet-pLKO-puro all-in-one vector (RRID: Addgene_21915) including a puromycin resistance cassette and a tet-responsive element for expression of shRNAs was established according to a publicly available protocol (46) using In-Fusion HD Cloning Kit (Clontech) (**Supplementary Table 3**). Vectors were amplified in Stellar Competent Cells (Clontech). Successful shRNA integration was verified by Sanger sequencing (primer: 5’-GGCAGGGATATTCACCATTATCGTTTCAGA-3’). Packaging plasmids pCMV-dR8.91 or psPAX2 (RRID: Addgene_12260), envelope plasmid pMD2.G (VSV-G) (RRID: Addgene_12259) and Tet-pLKO-puro all-in-one vectors harboring shRNAs against *RRM2* (shRRM2) or a non-targeting control shRNA (shControl) were transfected in HEK293T. Lentiviral particles were yielded by filtering the supernatant and infected in EwS cell lines A-673, ES7, and TC-71, followed by resistant cell selection with 1 µg/ml puromycin (InVivoGen, USA). From puromycin-selected cells single cell clones were established and those with sufficient knockdown levels (approximately onto 30% remaining expression) were used for further analyses. Knockdown induction was achieved by adding 0.1 µg/ml Dox in cell culture media every 48h. Established cell lines were designated as cell line/TR/shRRM2_1, cell line/TR/shRRM2_3, cell line/TR/shControl.

### Proliferation assays

2–3×10^4^ cells per well (depending on the cell line) were seeded on a 6-well plate in three technical replicates. 0.1 µg/ml Dox was added to media every 48h for knockdown induction. After harvesting cells including the supernatant, vital and dead cells were counted with a cell counter (Countess™ II Automated Cell Counter, Invitrogen) using Trypan-Blue exclusion method (Sigma-Aldrich).

### Clonogenic growth assays

Cells were seeded at a density of 0.5–5×10^3^ per well (depending on the cell line) on a 6-well plate in three technical replicates and grown for 9–12 days. For genetic target inhibition 0.1 µg/ml Dox was added to media every 48h. For pharmacological target inhibition cells were treated with either drugs in indicated doses or equimolar vehicle (DMSO). Media were changed and drugs renewed every 72h. Colonies were stained with crystal violet solution (Sigma-Aldrich). The colony area and intensity were calculated with the ImageJ Plugin *Colony area* (47).

### Murine xenograft model

2–2.5×10^6^ cells resuspended in a mix of 1:1 PBS and Geltrex Basement Membrane Mix (Thermo Fisher Scientific) were subcutaneously injected in the right flank of 10–12 weeks old NOD/Scid/gamma (NSG) mice. Tumor growth was assessed by measuring tumor size in two-dimension with a caliper every 2–3 day. The tumor volume was estimated by the formula length×(width)2/2. Once subcutaneous tumors became palpable (approx. 5 mm in diameter), animals were randomized in treatment group or control group. For induction of the *RRM2* knockdown 2 mg/ml Dox (BelaDox, Bela-pharm, Germany) was added in drinking water containing 5% sucrose (Sigma-Aldrich) for *ad libitum* uptake (Dox (+)), whereas the control group received sucrose alone (Dox (–)). For pharmacological RRM2 inhibition the treatment group received intraperitoneal injection of triapine 30 mg/kg every second day, while the control group received vehicle (DMSO) alone. The endpoint was set at reaching either an estimated tumor volume of 1,500 mm3 or average tumor diameter of 1.5 cm where animals were sacrificed by cervical dislocation. Any animals were excluded in case no subcutaneous tumor growth was observed. Animal experiments were approved by the government of Upper Bavaria and performed in accordance with ARRIVE guidelines, recommendations of the European Community (86/609/EEC), and United Kingdom Coordinating Committee on Cancer Research (UKCCCR) guidelines for the welfare and use of animals in cancer research.

### Cell viability assays and dose response assessment

1.5–5×10^3^ cells per well (depending on the cell line) were seeded on a 96-well plate and treated with compounds in a dose range for which clinically achievable doses were taken into account if applicable. Assays were performed in a total volume of 100 µl on at least three technical replicates. Cell viability was assessed 84h after the treatment start with Resazurin cell viability assays (16 µg/ml, Sigma-Aldrich). Dose response curves were simulated by nonlinear regression models and IC50 values were calculated using PRISM 9 (GraphPad Software Inc., CA, USA) by normalizing to the respective controls (vehicle alone).

### Assessment of drug interaction and combination efficiency

1.5–5×10^3^ cells per well (depending on the cell line) were seeded on a 96-well plate in triplicate and treated with various drug combinations either in constant dose ratios or in three serial doses (4×4 matrices). Cell viability was assessed 84h after the treatment start with Resazurin cell viability assays (16 µg/ml, Sigma-Aldrich) normalized to the respective controls (vehicle alone). The combination efficiency was analyzed using the Chou-Talalay method (48) with CompuSyn, in which the combination index (CI) was calculated based on the dose-effect property for single drugs and their combinations. Dose-effect curves were linearized with the median-effect-plot for each single drug and combinations, and doses which cause the equivalent effect (reduction of cell viability) by the single drugs or combinations were calculated. CI is given by the formula: CI= (D)1/(Dx)_1_+(D)_2_/(Dx)_2_. (Dx)_1_, doses of drug 1 alone which shows X% inhibition; (Dx)_2_, doses of drug 2 alone which shows X% inhibition; (D)_1_, dose portion of drug 1 in combination with drug 2, which shows X% inhibition; (D)_2_, dose portion of drug 2 in combination with drug 1, which shows X% inhibition. The combination efficiency was interpreted as: CI value < 1 indicative of synergistic, CI = 1 additive, and CI > 1 antagonistic. The estimated drug combination efficiency was calculated by SynergyFinder 2.0 (49) based on ZIP reference model. ZIP synergy score > 10, likely to be synergistic; between -10 and 10, likely to be additive; < –10, likely to be antagonistic.

### Establishment of drug-resistant EwS cells

Doxorubicin-, gemcitabine-, or triapine-resistant EwS cells (DR, GR, TR, respectively) were established by culturing parental cells with serially increased concentration of each drug. Parental cells were treated with each drug from initial concentrations equivalent to IC_50_ values as assessed by Resazurin cell viability assays. Cells were deemed to have successfully adapted to the increment once they started to constantly regrow. The degree of drug resistance was assessed by IC_50_ values as determined by Resazurin cell viability assays.

### Histology, immunohistochemistry (IHC), and evaluation of immunoreactivity

Formalin-fixed and paraffin-embedded (FFPE) tissue sections were routinely processed and stained with hematoxylin and eosin for histological assessment including tissue structure, cellular morphology, mitosis, and cell death. For the detection of γH2A.X (phosphor-S139) 4 μm FFPE tissue sections were cut followed by an antigen retrieval by heat treatment with Target Retrieval Solution (S1699, Agilent Technologies, Germany). Slides were incubated with a monoclonal anti-γH2A.X (phosphor-S139) primary antibody (rabbit, 1:8,000, ab81299, Abcam, UK) for 60 min at room temperature, followed by incubation with a monoclonal secondary horseradish peroxidase (HRP)-coupled horse-anti-rabbit antibody (ImmPRESS Reagent Kit, MP-7401, Vector Laboratories, Germany) with AEC+ (K346, Agilent Technologies, USA) as chromogen, counterstained by hematoxylin Gill’s formula (H-3401, Vector Laboratories, Germany). For detection of cleaved caspase 3, antigen retrieval was carried out by heat treatment with Target Retrieval Solution Citrate pH6 (S2369, Agilent Technologies). Slides were incubated with a polyclonal cleaved caspase 3 primary antibody (rabbit, 1:100; 9661, Cell Signaling, Frankfurt am Main, Germany) for 60 min at room temperature followed by incubation with a monoclonal secondary horseradish peroxidase (HRP)-coupled horse-anti-rabbit antibody with AEC+ as chromogen, counterstained by hematoxylin Gill’s formula. For detection of RRM2, antigen retrieval was performed by heat treatment with ProTaqs IV Antigen-Enhancer (Quartett, 401602392). Slides were incubated with a polyclonal anti-RRM2 primary antibody (rabbit, 1:500, atlas antibodies, HPA056994) for 60 min at room temperature followed by incubation with a monoclonal secondary horseradish peroxidase (HRP)-coupled horse-anti-rabbit antibody with AEC+ as chromogen, counterstained by hematoxylin Gill’s formula (H-3401, Vector Laboratories, Germany). Slides were scanned using a Nanozoomer-SQ Digital Slide Scanner (Hamamatsu Photonics K.K.) and visualized with NDP.view2 image viewing software (Hamamatsu Photonics K.K.).

For assessment of RRM2 expression levels, immunoreactivity was semi-quantified as previously described in analogy to the hormone receptor scoring system Immune Reactive Score (IRS) with slight modifications to account for a 5-tier grading of positive tumor area (10, 50). First, the percentage of immunoreactive tumor cells was graded as follows; grade 0 = 0−19%, grade 1 = 20−39%, grade 2 = 40−59%, grade 3 = 60−79% and grade 4 = 80−100%. Second, the relative intensity of immunoreactivity was classified as follows; grade 0 = none, grade 1 = low, grade 2 = moderate and grade 3 = strong. IRS was then given as the product of both grades. For assessment of cleaved caspase-3 (CC3) and γH2A.X, immunoreactive cells were counted under the microscope in ten high power fields.

### Transcriptome profiling, gene set enrichment analysis (GSEA), and gene ontology (GO) analysis

For transcriptome profiling experiments, 1.0–2.0×10^5^ A-673/TR/shRRM2_1, A-673/TR/shRRM2_3, A-673/TR/shControl, ES7/TR/shRRM2_1, ES7/TR/shRRM2_3 and ES7/TR/shControl were seeded in T-25 flasks, and treated with 0.1 µg/ml Dox for 72h (A-673) and 96h (ES7). For pharmacological RRM2 inhibition A-673 and ES7 were treated with either triapine (IC_50_ for A-673 0.44 µM or for ES7 0.65 µM) or equimolar vehicle (DMSO) for 72h. Total RNA was extracted using the NucleoSpin RNA kit according to the manufacturer’s instruction (Macherey-Nagel, Germany). Samples with RNA integrity numbers (RIN) >9 were hybridized to Human Affymetrix Clariom D microarrays at IMGM Laboratories (Munich, Germany). Data were quantile-normalized with Transcriptome Analysis Console (v4.0; Thermo Fisher Scientific) using the SST-RMA algorithm as previously described (51). For gene annotation the Affymetrix library for Clariom D Array (version 2, human) was employed.

For identification of differentially expressed genes (DEGs) with consistent and significant fold changes (FCs) across shRNAs and cell lines, genes with log2 transformed gene expression values lower than that of *ERG* (mean log2 expression around 6.0), which is virtually not expressed in EWSR1-FLI1 positive EwS cell lines (52) were excluded. The FCs in shControl or vehicle, and two specific shRNAs or triapine treatment samples were individually calculated for each cell line. The FCs in shControl or vehicle samples were subtracted from those of shRRM2 or triapine treatment samples, respectively, yielding the cell line specific FCs for two specific shRNAs or triapine treatment. To integrate FCs across shRNAs or cell lines average FCs were used for further analyses. For downstream analyses, those genes with a minimum absolute log2 FC of 0.5 were included. To identify enriched gene sets, genes were ranked by the FC values, and a pre-ranked GSEA (MSigDB v7.0, c2.cpg.all) with 1,000 permutations was performed. To analyze enriched gene sets and their correlation upon RRM2 silencing, Weighted Gene Correlation Network Analysis (WGCNA) was performed (53) using Gene Ontology (GO) biological processes terms from MSigDB (c5.all.v7.0.symbols.gmt). Enriched GO terms were filtered for statistical significance (adjusted *P*<0.05) and a normalized enrichment score |(NES)|>1.5 (10,000 permutations). The constructed correlation network was visualized using Cytoscape (54).

For GO enrichment analysis under the condition of *RRM2* silencing or pharmacological inhibition in each cell line, commonly regulated genes in both cell lines and two experimental settings were extracted using Draw Venn Diagram (Van de Peer Lab, http://bioinformatics.psb.ugent.be/webtools/Venn/). The commonly regulated genes were then interrogated for overrepresentation of biological processes using GO enrichment analysis (55–57). For gene co-expression analysis, the gene expression correlation between *RRM2* and other genes from 166 EwS tumors was estimated by calculating Pearson correlation coefficients. Those genes with |r_Pearson_| > 0.5 were further subjected to GO enrichment analysis. Heat maps were created by Morpheus (https://software.broadinstitute.org/morpheus) using the Pearson correlation coefficients between *RRM2* and other genes. Gene expression data were deposited at the GEO (accession codes: GSE166415 and GSE166419).

### Human tissue samples and ethics approval

Human tissue samples were retrieved from the tissue archives of the Institute of Pathology of the LMU Munich (Germany) or the Gerhard-Domagk Institute of Pathology of the University of Münster (Germany) upon approval of the institutional review board. All patients provided informed consent. Tissue-microarrays (TMAs) were stained and analyzed with approval of the ethics committee of the LMU Munich (approval no. 550-16 UE).

### Analysis of copy number variation and promoter methylation in primary EwS

For analysis of genomic copy numbers, publicly available DNA copy number data for EwS tumors (58) and corresponding RNA expression data (GSE34620 and GSE37371, *n*=32) were downloaded from the ‘soft tissue cancer – Ewing sarcoma – FR’ project from the International Cancer Genome Consortium (ICGC) Data Portal and Gene Expression Omnibus (GEO) of the NCBI, respectively. Segment mean values for *RRM2* locus were obtained using Visual Basic for Applications (VBA). The correlation between the segment mean values and log2 transformed *RRM2* expression was analyzed using PRISM 9 (GraphPad Software Inc., CA, USA). For analysis of CpG methylation on the *RRM2* promoter region, publicly available data on CpG methylation in 40 EwS tumors (GSE88826) (59) and corresponding RNA expression data (GSE34620) were downloaded from GEO. The correlation of the methylation rates and *RRM2* expression on 5 CpG sites (CpG1 hg19: chr2:10120351; CpG2 hg19: chr2:10122310; CpG3 hg19: chr2:10122312; CpG4 hg19: chr2:10122323; CpG5 hg19: chr2:10122335) in each sample (*n*=40) was analyzed using PRISM 9 (GraphPad Software Inc., CA, USA).

### Statistical analyses

Statistical data analyses were performed using PRISM 9 (GraphPad Software Inc., CA, USA) on the raw data. For hypothesis tests for two groups a two-sided Mann-Whitney test was used if not otherwise specified in the figure legends. A Fisher exact probability test was employed for contingency data sets. To analyze bivariate correlations, Pearson correlation coefficients were calculated and analyzed with PRISM 9. Data are presented as scatter-bar-plots with horizontal bars indicating means and whiskers the standard error of the mean (SEM), if not otherwise specified in the figure legends. The sample size for *in vitro* experiments was chosen empirically. The sample size for *in vivo* experiments was predetermined using power calculations with *β* = 0.8 and *α* <0.05 based on preliminary data and in compliance with the 3R system (replacement, reduction, refinement). For survival analysis, overall survival for clinical data sets or event-free survival for *in vivo* experiments were described by Kaplan-Meier curves, and survival functions between the groups were analyzed by a log-rank test or a Mantel-Haenszel test. *P*-values < 0.05 were considered as statistically significant. *P*-values were calculated from two-sided statistical tests, if not otherwise specified in the figure legends.

### Data and code availability

Original microarray data used in this study were deposited at the National Center for Biotechnology Information (NCBI) GEO under accession numbers GSE166415 and GSE166419. Custom code is available from the corresponding author upon reasonable request.

## LEGENDS TO SUPPLEMENTARY TABLES

**Supplementary Table 1 Overexpressed genes in EwS compared to normal tissues and their prognostic relevance.**

**Supplementary Table 2 Commonly regulated genes upon *RRM2* silencing and pharmacological inhibition by triapine.**

**Supplementary Table 3 Oligonucleotide sequences.**

## LEGENDS TO SUPPLEMENTARY FIGURES

**Supplementary Fig. 1.**
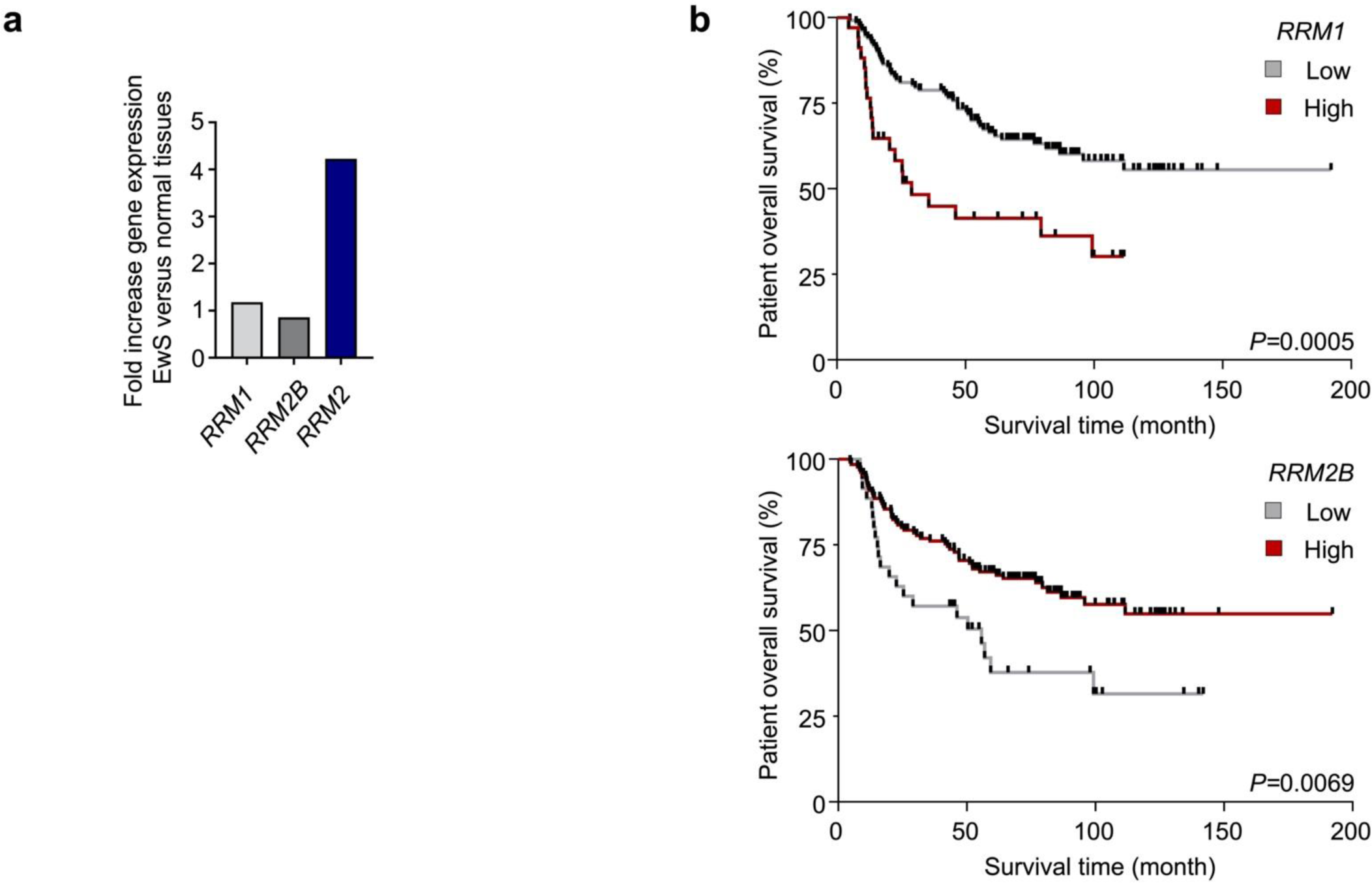
*RRM1* and *RRM2B* expression in EwS and normal tissues, and their association with overall survival in 166 EwS patients. **a)** Analysis of *RRM1* and *RRM2B* mRNA expression levels in 50 EwS primary tumors compared to 929 normal tissues samples from 71 tissue types. Data are shown by fold increase normalized to expression values of normal tissues. **b)** Analysis of overall survival time of 166 EwS patients for *RRM1* and *RRM2B* mRNA expression. *P*-values were determined in Kaplan-Meier analyses using a Mantel-Haenszel test.

**Supplementary Fig. 2.**
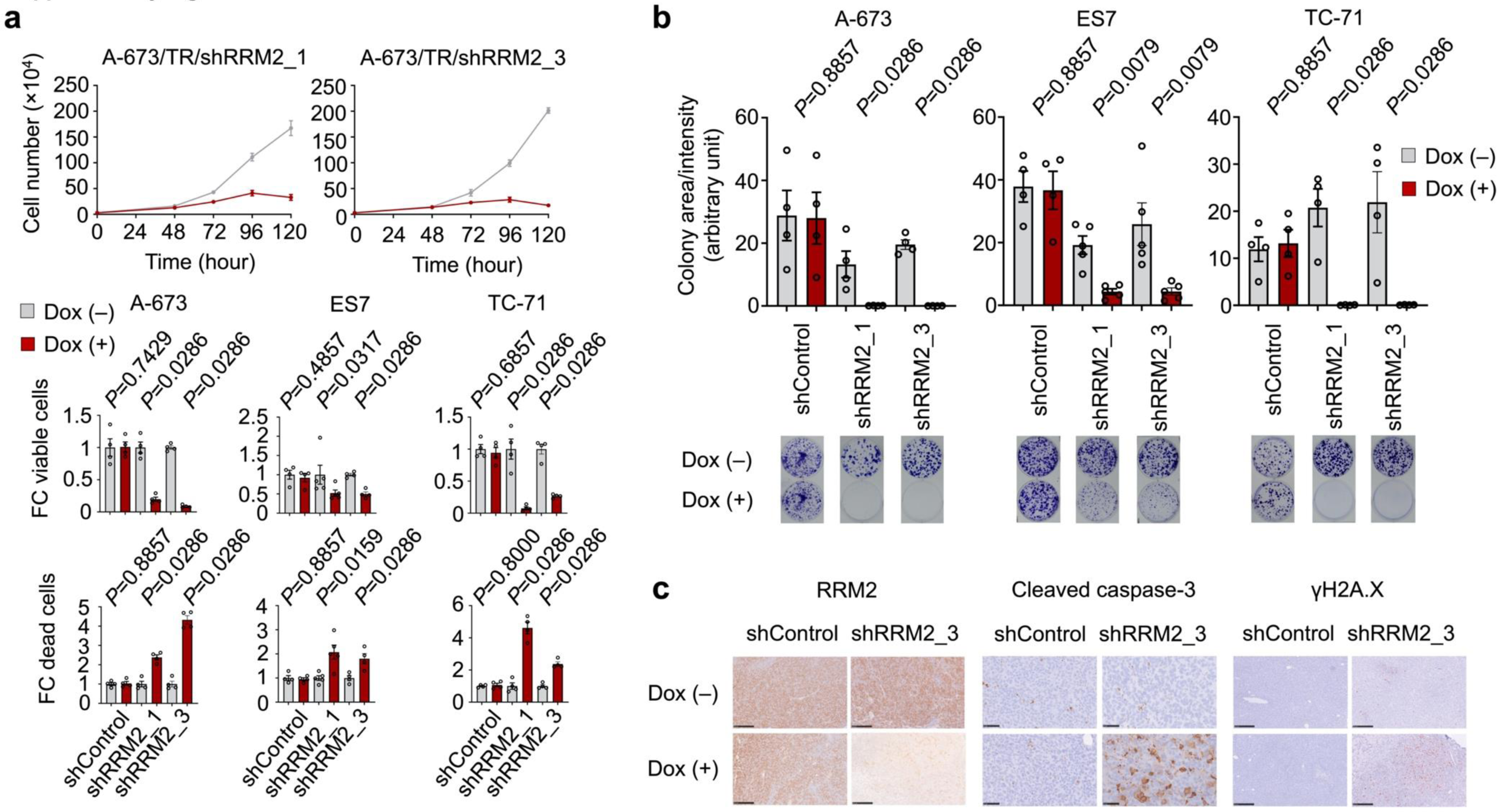
*RRM2* silencing inhibits cell proliferation and clonogenic growth in EwS *in vitro*. **a)** Analysis of proliferation assays upon shRNA-mediated *RRM2* silencing in EwS cell lines. Upper: Cell proliferation over 120h upon *RRM2* silencing in A-673. Viable cells upon *RRM2* silencing (middle) and dead cells (lower) in EwS cell lines (A-673, ES7, TC-71) harboring Dox-inducible shRRM2 constructs or non-targeting shRNA (shControl). Values are normalized to shControl. Horizontal bars represent means and whiskers SEM. FC, fold change. Two-sided Mann-Whitney test. **b)** Analysis of clonogenic growth upon shRNA-mediated *RRM2* silencing in EwS cell lines (A-673, ES7, TC-71) harboring Dox-inducible shRRM2 constructs or non-targeting shRNA (shControl). Horizontal bars represent means and whiskers SEM. Two-sided Mann-Whitney test at the experimental endpoint.

**Supplementary Fig. 3.**
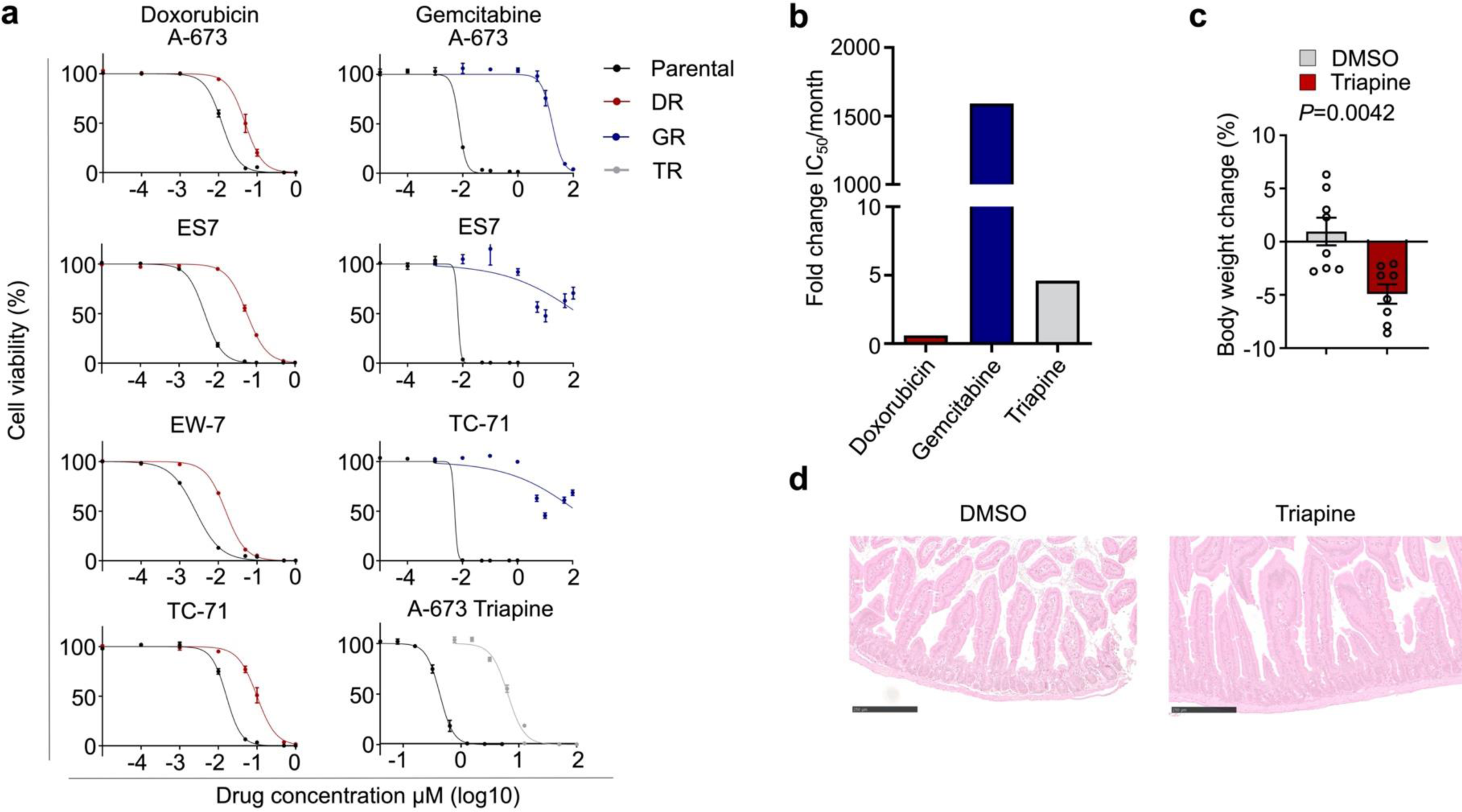
Development of chemoresistance in EwS cell lines and adverse effects of triapine treatment *in vivo*. The EwS cell lines were subjected to serially ascending doses of either doxorubicin (A-673, ES7, EW-7, TC-71), gemcitabine (A-673, ES7, TC-71) or triapine (A-673). **a)** Comparative drug response analysis for doxorubicin or gemcitabine in parental, doxorubicin-resistant (DR) or gemcitabine-resistant (GR) cells assessed by Resazurin cell viability assays. **b)** Drug-resistance developing rate for doxorubicin, gemcitabine or triapine in A-673. Data are shown by fold change in IC_50_ normalized to those of the parental cells divided by time for stably acquiring drug resistance. **c)** Change of body weight upon triapine or vehicle treatment at the experimental endpoint. Data are shown by body weight change before and after treatment normalized to the baseline (initial values before starting treatment). **d)** Histological assessment of adverse effects in the intestine. Hematoxylin and Eosin staining. Scale bar=250 µm.

**Supplementary Fig. 4.**
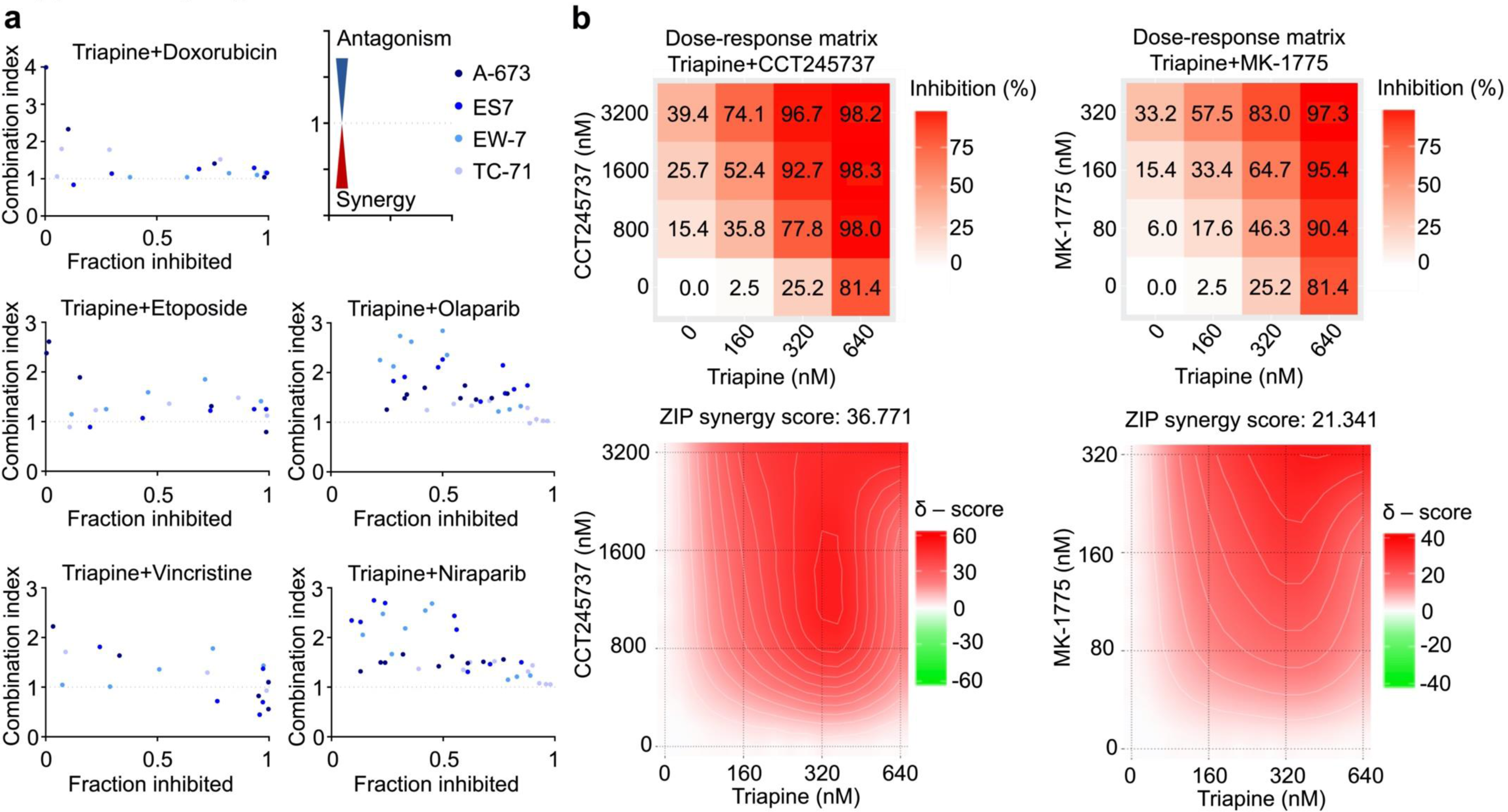
Drug interaction and combination efficiency of triapine with chemotherapeutics, PARP inhibitors, CHEK1 inhibitor or WEE1 inhibitor. **a)** Drug interaction and combination efficiency analysis between triapine and doxorubicin, etoposide, vincristine, olaparib or niraparib in 4 EwS cell lines (A-673, ES7, EW-7, TC-71) assessed by combination index. CI value < 1 indicative of synergistic, CI = 1 additive, and CI > 1 antagonistic **b)** Drug interaction and combination efficiency estimation between triapine and CHEK1 inhibitor (CCT245737) or WEE1 inhibitor (MK-1775) in A-673 EwS cell line assessed by SynergyFinder 2.0. Upper: Dose response matrix. Lower: Synergy distribution. ZIP synergy score > 10, likely to be synergistic; between -10 and 10, likely to be additive; < –10, likely to be antagonistic.

**Supplementary Fig. 5.**
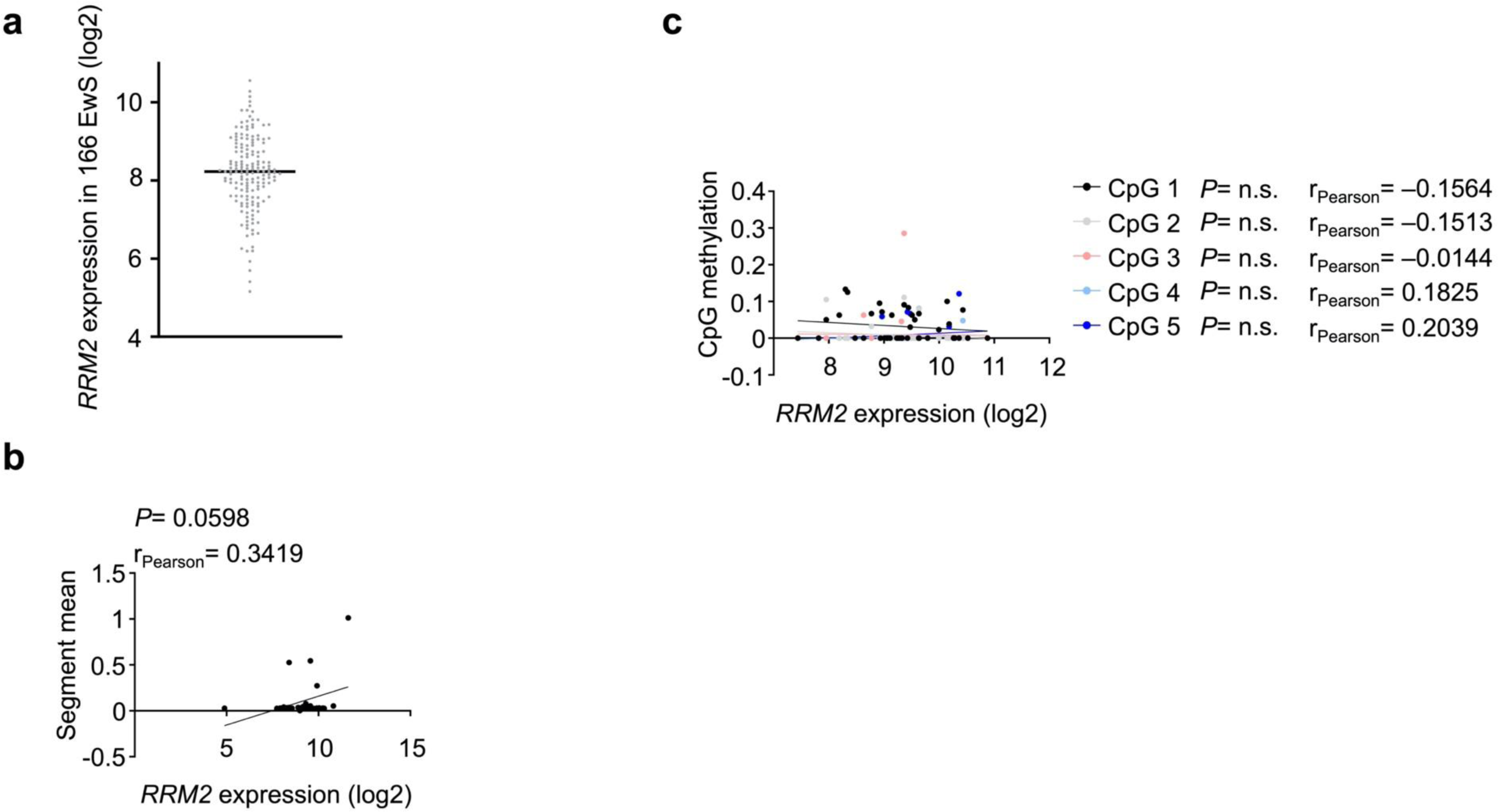
Distribution of *RRM2* expression in 166 EwS, copy number variations (CNVs) and promoter methylation status at the *RRM2* locus in EwS. **a)** Distribution analysis of *RRM2* mRNA expression in 166 EwS patients. Each dot represents individual *RRM2* expression. **b)** Correlation analysis of CNVs at the *RRM2* locus with *RRM2* mRNA expression levels in primary EwS tumors (n=32). The solid line indicates a trend line estimated by a simple linear regression model. **b)** Correlation analysis of promoter methylation on five CpG sites with *RRM2* expression levels in primary EwS tumors (n=40). The solid lines indicate trend lines estimated by a simple linear regression model.

